# Explicit integration of dispersal-related metrics improves predictions of SDM in predatory arthropods

**DOI:** 10.1101/2020.06.05.136044

**Authors:** Monsimet Jérémy, Devineau Olivier, Pétillon Julien, Lafage Denis

**Affiliations:** Department of Forestry and Wildlife management, Inland Norway University of Applied Sciences, Campus Evenstad, Koppang, Norway; UMR CNRS 6553 ECOBIO, Université de Rennes, Rennes, France; Department of Environmental and Life Sciences/Biology, Karlstad University, Karlstad, Sweden

**Keywords:** Fishing spiders, Pisauridae, Climate change, Dispersal limitation, Hybrid SDM

## Abstract

Fishing spiders (*Dolomedes spp.*) make an interesting model to predict the impact of global changes because they are generalist, opportunistic predators, whose distribution is driven mostly by abiotic factors. Yet, the two European species are expected to react differently to forthcoming environmental changes, because of habitat specialization and initial range. We used an original combination of habitat and dispersal data to revisit these predictions under various climatic scenarios. We used the future range of suitable habitat, predicted with habitat variables only, as a base layer to further predict the range or reachable habitat by accounting for both dispersal ability and landscape connectivity. Our results confirm the northward shift in range and indicate that the area of co-occurrences should also increase. However, reachable habitat should expand less than suitable habitat, especially when accounting for landscape connectivity. In addition, the potential range expansion was further limited for the red-listed *D. plantarius*, which is more habitat-specialist and has a lower ability to disperse. This study highlights the importance of looking beyond habitat variables to produce more accurate predictions for the future of arthropods populations.

## Introduction

Climate change, which is now threatening all ecosystems worldwide (Bellard et al. 2012), is a multi-factor problem that goes beyond raising temperatures only (Pereira et al. 2010, Garcia et al. 2014). Tackling this complexity requires that ecologists obtain realistic predictions of how species distributions will change in response to global change. In recent years, species distribution models (SDMs) proved to be an important tool to for this. SDMs are particularly useful to predict geographic distributions by correlating species occupancy to environmental variables (Miller 2010). Applications include conservation planning (Guisan et al. 2013), potential invasion range (Bellard et al. 2013), or forecasting in time (Hijmans and Graham 2006). SDMs were successfully applied to a large variety of terrestrial (see Hao et al. 2019 for a review) and marine organisms (see Melo-Merino et al. 2020 for a review).

The accuracy of predictions produced by SDMs varies from algorithm to algorithm, even when considering that the MaxENT algorithm is most often used (Qiao et al. 2015). This variation in accuracy can be alleviated with ensemble models, which combine algorithms and produce consensual predictions (Arauja and New 2007, Thuiller 2004). Of course, input data also influence the predictions (Thuiller et al. 2019), and while most SDMs use only climatic variables, including other variables such as land-use might improve predictions (Titeux et al. 2016). In order to make projections in time, it is fundamental to carefully select the right climatic scenario (Thuiller et al. 2019). Right now, the ones produced and updated by the Intergovernmental Panel on Climate Change (IPCC 2007) are the most widely recognized and used climatic scenarios.

SDMs assume that the species and its environment are at equilibrium (Guisan and Thuiller 2005), so that all suitable locations are occupied. SDMs also assume that the ecological niche is stable, i.e. that the same factors limit the species in space and time (Richmond et al. 2010). Under these assumptions, SDMs are used to define habitat suitability, which is the range of physical locations where one species can live (Kearney 2006). However, a properly constructed and calibrated SDM can provide information about the specie’s realized niche, ie a combination of habitat with other biotic and abiotic factors (Guisan and Thuiller 2005, Soberon and Peterson 2005).The gold standard of SDMs would be fully mechanistic models which were used, for example to study seed dispersal in birds (Merow et al 2011) or population dynamics and evolution of dispersal trait (Bocedi et al. 2017). However, these models are very data-demanding, and simpler hybrid mechanistic-correlative models are often more suitable for less well-studied taxa. In particular these hybrid models allow including active biological processes such as dispersal (Briscoe et al. 2019). Examples include making predictions under full /no dispersal (Thuiller et al. 2009) or using a buffer of dispersal around each presence (Mammola and Isaia 2017).

As generalist predators, spiders are relatively independent of a specific prey community, and their assemblage and distribution is mostly influenced by habitat and land use (Lafage et al. 2015), which makes them good study cases for SDMs. Fennoscandia is a potential climatic refugium for spider populations against the current global warming (Leroy et al. 2014). Refugia can mitigate the effects of climate change by providing suitable conditions for species persistence through time (Keppel and Wardell-Johnson 2012). *Dolomedes plantarius* could presumably use Fennoscandia as a refugium, but the ability of the species to effectively spread northward has not been accounted for in previous predictions (Leroy et al. 2013, 2014). Moreover, fishing spiders are threatened by the decrease of range and quality of their wetland and fenland habitats, which are declining globally (Finlayson et al. 2019). The other European fishing spider, *Dolomedes fimbriatus*, also occurs in Fennoscandia. Co-occurrence of both *Dolomedes*, was considered impossible due to different habitat requirements (van Helsdingen 1993). Syntopy is possible though, as the two species can live close to each other (Duffey 2012), for example around the same lake (Ivanov et al. 2017), or in the ecotone habitat between bogs and ponds (Holec 2000). *D. fimbriatus* has a larger ecological niche: the species is more drought and shade tolerant (Duffey 1995), and is less sensitive to water quality (Duffey, 2012). Consequently, *D. fimbriatus* could become a competitor to *D. plantarius* in syntopic sites if global change brings more frequent drought events

Here, we compare the potential range spread of *D. plantarius* and *D. fimbriatus*, and their ability to use Fennoscandia as a refugium. We aim to provide more conservative predictions for Fennoscandia than previously predicted at the European scale by Leroy et al (2013). To do so, we developed hybrid species distribution models including climate and land-use variables, as well as dispersal and landscape connectivity (figure 1). We expected that:

**Figure 1:**
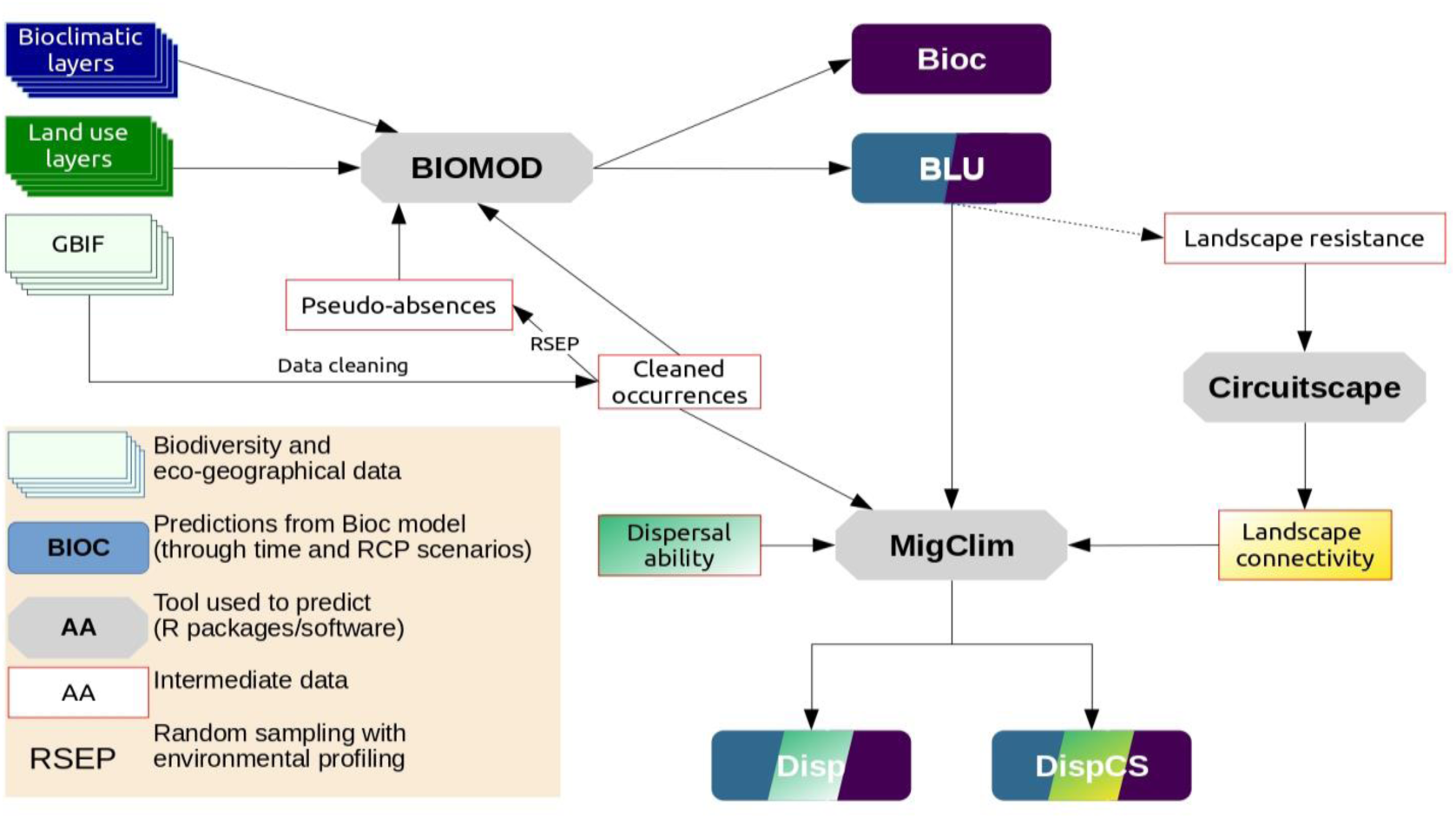
Flowchart of the framework used to study the future distribution of the two European fishing spiders (Bioc: bioclimatic only model, BLU: bioclimatic and land use model, Disp: dispersal model, DispCS: dispersal and landscape connectivity model).

1. The distribution of both fishing spiders should expand northward (Parmesan and Yohe 2003, Parmesan 2006). A larger expansion is expected under more intense climate change.
2. Since *D. fimbriatus* is a habitat generalist, the range of habitat it can reach should be larger and occupied faster, than for *D. plantarius* (Hill et al. 1999).
3. The area of sympatry between the two species should increase with the range expansion of the two species.

## Methods

### Occurrence data

We downloaded records of presence for both spider species from the GBIF (GBIF: The Global Biodiversity Information Facility 2019) via the rgbif package (citations for R packages are provided in Supplementary material, Appendix 1) in R (R Core Team 2019). The GBIF database gathers volunteer-based naturalist observations (Supplementary material Appendix 2), which often require a quality check. We used the package CoordinateCleaner (Supplementary material Appendix 1) to remove null or duplicate coordinates, and to flag the records requiring a subjective decision, such as ol records or records located in urban areas, or at the centroid of a county. Urban records were not necessarily false presence, and we used aerial photography (ESRI 2009) accessed with packages leaflet and mapedit (Supplementary material Appendix 1) to decide whether to keep these records or not. We visually checked, for instance, if a record was not in a recently modified areas in a city. Some records suggesting co-occurrence of the two species were checked in the field during summer 2018 and 2019 (25 locations, including four actually syntopic locations). We retained 775 records for *Dolomedes fimbriatus* and 181 records for *Dolomedes plantarius* (Figure 2), reflecting the GBIF data available until October 2019 in Fennoscandia. When several records fell in the same raster cell, we kept only one.

**Figure 2:**
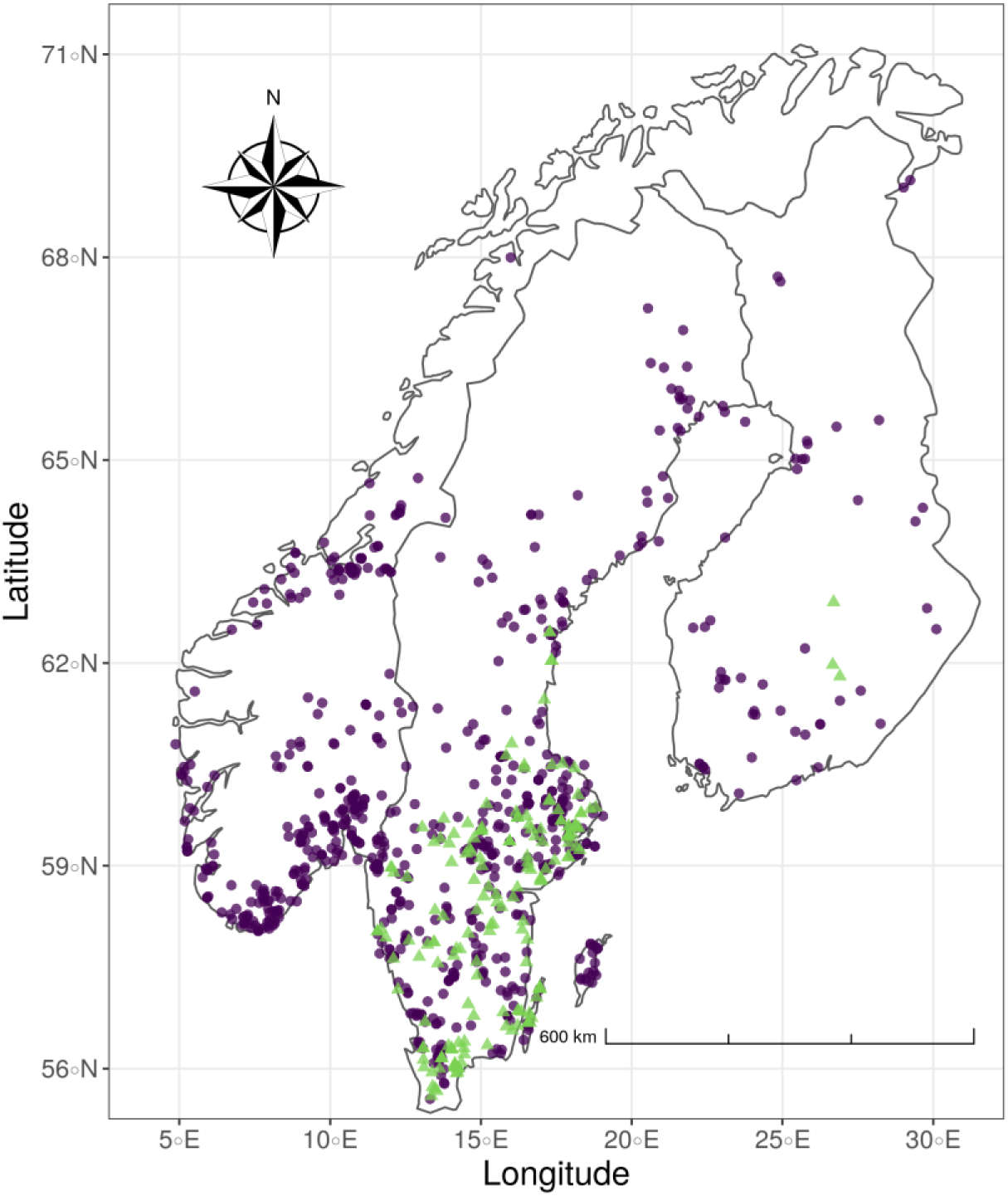
*Dolomedes plantarius* (green triangles) and *Dolomedes fimbriatus* (purple dots) records in Fennoscandia as of October 2019. Data were extracted from the GBIF database and supplemented by field samplings.

### Species distribution modelling

#### Predictor variables

For the climatic component of the ecological niche, we included variables which were biologically relevant for spiders, and not too correlated (Braunisch et al. 2013). Using a correlation coefficient threshold of 0.7 (Dormann et al. 2012), we selected mean and maximum annual temperature, mean diurnal temperature range, mean temperature of the wettest quarter, and annual precipitation, which we extracted from the WorldClim database (Fick and Hijmans 2017) at a spatial resolution of 30 arc-seconds (Supplementary material Appendix 4, Tab. A1).

To predict the future distribution of *Dolomedes* spiders in Fennoscandia, we used IPCC projections for 2050 and 2070, under multi-factors “representative concentration pathways” (RCP) 4.5 and 8.5 (van Vuuren et al. 2011). RCP4.5 corresponds to medium-low greenhouse gas emissions and air pollution, whereas RCP8.5 considers high greenhouse gas emission, medium air pollution, and an increase in carbon dioxide (van Vuuren et al. 2011). We downloaded these climatic projections from Wordclim (Hijmans et al. 2005) at a spatial resolution of 30 arc-seconds.

For the habitat component of the ecological niche, we integrated information on ground wetness, which is an important community driver for the semi-aquatic fishing spiders (Lafage et al. 2015, Lafage and Pétillon 2016). We also incorporated forest and grassland density, because the presence of fishing spiders seems to be influenced by the surrounding landscape (unpublished data). We downloaded the corresponding geographic layers from the Copernicus Land Monitoring Service at 100-metres resolution (EEA 2018), and upscaled them to 30 arc-seconds resolution to match the bioclimatic data. The forest layer represents the density of the tree cover (from 0 to 100 %) in 2015. The ‘Water and Wetness’ layer represents the occurrence of wet surfaces from 2009 to 2015, using a water and wetness probability index, indicating the degree of physical wetness, independently of the vegetation cover. Finally, the grassland layer represents the percentage of grassland per pixel. We estimated the change in land use between current and future times with a model which harmonises scenarios from different integrated assessment models, namely MESSAGE for RCP8.5 and GCAM for RCP4.5 (Hurtt et al. 2011).

#### Calibration area and pseudo-absences

To use presence-absence models with the presence-only GBIF data, we used a random sampling procedure with environmental profiling (RSEP; Senay et al. 2013), which creates a background of absence records for each algorithm. We generated the pseudo-absences in a different calibration area for each species. *D. plantarius* is a lowland species, so its calibration area was at low altitude <1000m. For *D. fimbriatus*, we excluded areas >1500m.

#### Model validation

Although there are many SDMs, none stands out as better than the others (Qiao et al. 2015). To improve the predictions, we therefore used an ensemble forecast approach, which combines several models weighted by their predictive accuracy (Buisson et al. 2010, Grenouillet et al. 2011).

Following recommendations in Barbet-Massin et al. (2012), we built our ensemble model with 10 runs of gradient boosting models (GBMs), generalized additive models (GAMs) and Maxent. We used 1000 pseudo-absences for the GBMs, and as many pseudo-absences as presences for the GAMs. We used 80% of the data for training the ensemble model and testing the single run of model, and 20% for validation. Each model was cross-validated with a 5-fold procedure in package biomod2 (Supplementary material Appendix 1), thus leading to 5 fits for each type of model and each pseudo-absences run. We then evaluated the predictive accuracy of individual models with the true skill statistic (TSS) and the area under the receiving operating curve (AUROC). The TSS metric represents the ratio of hit rate to false alarm rate and varies from -1 to +1 (Allouche et al. 2006). We used a threshold of TSS = 0.4 to include models into the ensemble forecast (Allouche et al. 2006). The AUROC is a measure of “separability”, which represents the true positive rates graphically against the true negative rates. Following Fawcett (Fawcett 2006), we retained models with AUC>0.7 for the ensemble model. Finally, we converted the probabilities of presence predicted by the ensemble model into a binary presence/absence, with a cut point based on predictions which maximized the TSS (Supplementary material Appendix 1). In package biomod2, the relative variable contribution is assessed based on the correlation between the prediction of a model including a given variable and the model where this variable was dropped.

We built one model with bioclimatic variables only (model Bioc), and one with bioclimatic and land-use variables (model BLU). We then included dispersal to predict the range of suitable, but unreachable habitat (model Disp). Finally, we accounted for landscape connectivity into model dispCS. The framework is summarized in figure 1 (additional details in Supplementary material Appendix 4, Tab. A1).

#### Including dispersal into SDM

Although they differ in their general dispersal ability, the two species of fishing spider disperse mostly through ballooning and rappelling, where they catch the wind with a thread of silk, and passively fly. Laboratory tests suggested that few individuals exhibit long-distance dispersal behaviour on the water surface (unpublished data). We recorded this behaviour only in *Dolomedes fimbriatus* through sailing (when spider raised its body and/or abdomen and/or the legs to catch the wind). However, juveniles of *D. fimbriatus* are generally found in the surrounding vegetation rather than on the water (Duffey 2012), which makes aquatic dispersion unlikely.

We modelled dispersal ability via the MigClim package (Supplementary material Appendix 1), based on the predicted map of the BLU model. For each species, the MigClim model evaluates if suitable cells of the raster could become accessible between current time and 2050/2070. The package uses a dispersal kernel, i.e., a vector of probabilities of dispersal, to simulate the dispersal of the species (Supplementary material Appendix 3, Tab. A1). We used an imperviousness map (EEA 2018) to locate areas where the species settlement is highly unlikely. Since both fishing spiders are water-dependent, impervious regions where the soil seals, are barrier to settlement. Part of the MigClim modelling process is random (Engler and Guisan 2009), so we replicated each model 30 times and model-averaged the estimates.

In experimental settings, aerial dispersal (ballooning) is usually characterized when the spider is observed tiptoeing in response to a controlled wind. However, not all tiptoeing spiders end up ballooning (Bonte et al. 2009, Lee et al. 2015). The distance covered by aerial dispersal is less than 5 kilometres on average and is not correlated with the duration of the tiptoeing behaviour (Reynolds et al. 2007). We parametrized the MigClim model with values from the literature on aerial dispersal distance in spiders (Thomas et al. 2003, Reynolds et al. 2007). We weighed these values by the proportion of individuals we observed rappelling in our laboratory experiments (Monsimet et al. in prep), namely, 76.6% of *D. fimbriatus* and 59% *D. plantarius*. For long-distance dispersal, we used the proportion of individuals observed ballooning (*D. fimbriatus*: 14%, *D. plantarius*: 2.9%) for 2019. We considered that the probability of a settlement was similar for both species. Also, we hypothesized that it takes two years for a newly colonized area to produce new propagules, based on the >2-year lifespan of spiders in Northern Europe (Duffey 2012).

#### Accounting for landscape connectivity

We used the Circuitscape software (Shah and McRae 2008) to predict the potential dispersal corridors that *Dolomedes* could use to colonize their suitable habitat. Circuit theory estimates multiple pathways based on the resistance and conductance of the landscape (McRae et al. 2008). We used the habitat suitability prediction map from our BLU model to define the resistance map used by Circuitscape. We transformed the estimates of habitat suitability according to recommendations in Keeley (Keeley et al. 2017; see also Supplementary material Appendix 3).

We used a “wall-to-wall” approach (Pelletier et al. 2014, Febbraro et al. 2019) which estimates the conductivity of the landscape from South to North, and from West to East. A consensus map was produced by multiplying the resistance layers of different directions. This consensus map was an estimation of the landscape connectivity for the two species. The consensus map was binarized by considering conductance higher than mean conductance plus standard deviation as corridors (Febbraro et al. 2019). Areas outside corridors were then considered as a barrier to short-distance dispersal in Migclim. Migclim was parametrized as for the model Disp but accounting for the landscape connectivity barrier to make predictions for model DispCS.

### Range expansion and geographic overlap in time

We compared suitable habitat predicted across species, models, and scenarios. To estimate the range expansion or reduction in the future, we used the biomod2 package in R. We compared the direction of the shift in suitable habitat by calculating the centre of gravity of the suitable range with the SDMTools package (Supplementary material Appendix 1). To estimate the overlap of suitable habitat range between species for each time/scenario combination, we used the Schoeners’ D overlap metric (Warren et al. 2008), which ranges from 0 for no overlap to 1 for full overlap (Rödder and Engler 2011). We estimated the suitable habitat range overlap and not the full niche overlap here. We calculated D with the ENMtools package (Supplementary material Appendix 1).

## Results

### Modelling and model validation

The predictive performance of both Bioc and BLU models was higher than the threshold with either the ROC (>0.7) or the TSS (>0.4) metric (Supplementary material Appendix 4, Tab. A2). The relative contribution of predictors was the same across models and species, with mean annual temperature the most important variable with a contribution higher than 60%. For Bioc, mean temperature of the warmest month was also important, with a higher contribution for *D. fimbriatus* than for *D. plantarius* (33% and 11%, respectively). Mean temperature of the wettest quarter, annual precipitation and mean diurnal range contributed less than 10% to both models. Forest and ground wetness contributed more than grassland in the BLU models, but their relative contribution was less than 16%.

### Range expansion and geographic overlap in time

The size of the predicted / projected range was similar for both Bioc and BLU models. However, range expansion was predicted to be more restricted when also acccounting for land use (BLU) than when considering only climatic variables (Bioc). Indeed, adding land use variables contracted the suitable habitat at the limit of the range. Suitable range was also smaller for RCP4.5 than for RCP8.5, with similar patterns in time, except for *D.fimbriatus* where the range was reduced in 2070 compared to current under model BLU (figure 3).

**Figure 3:**
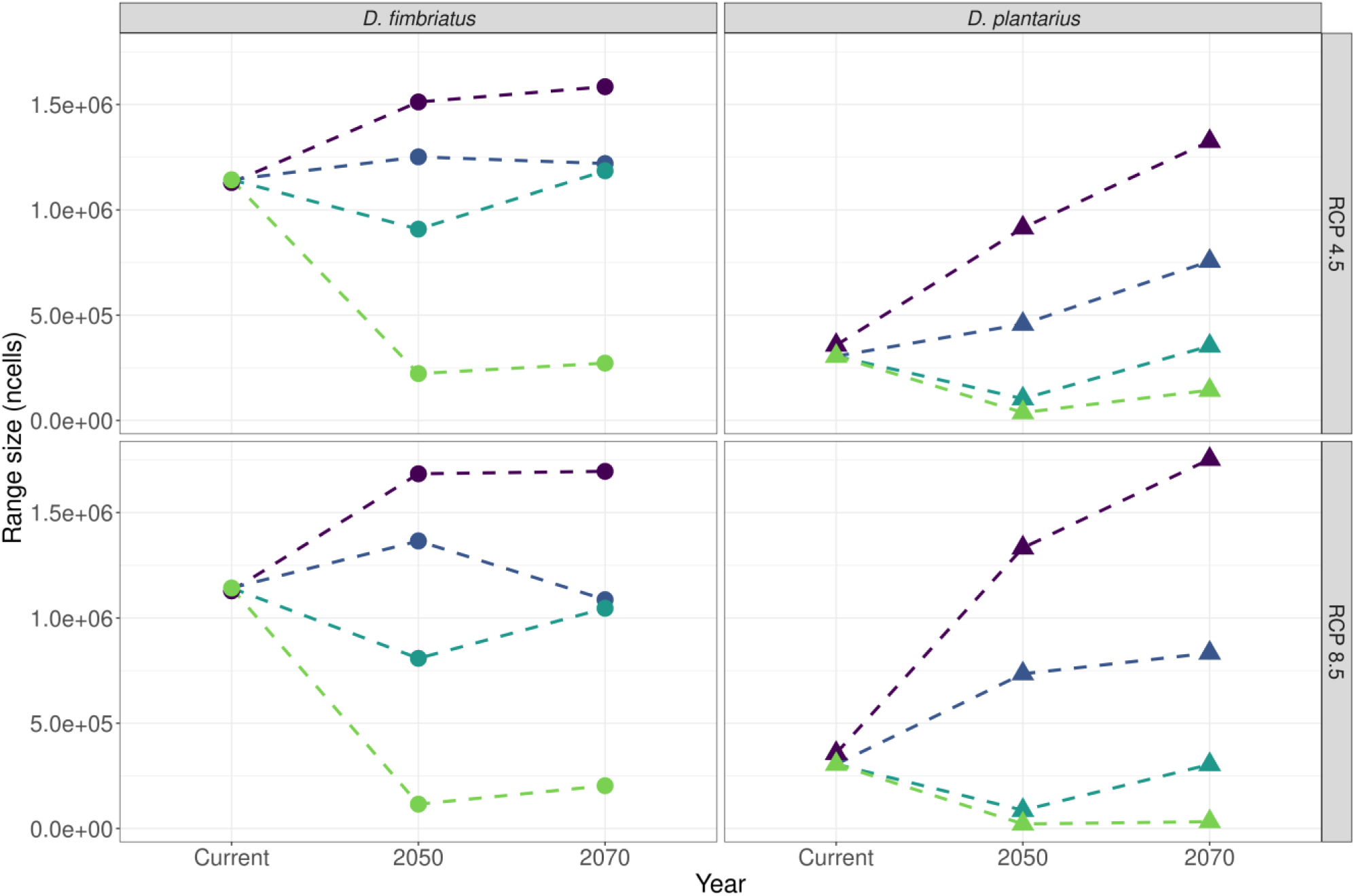
Range size in number of cells of suitable habitat predicted by the different SDMs in time per species and scenarios. (dark purple: Bioc model: bioclimatic variables only; dark blue: BLU model, bioclimatic + land use; Turquoise: Disp model with dispersal; green: DispCS model: dispersal and landscape connectivity).

Under RCP4.5 scenario, the suitable range was predicted to increase for both species in 2070 with the BLU model (14% for *D. fimbriatus* and 161% for *D. plantarius*). With model Disp, the range should decrease in 2050 for *D. fimbriatus* (20% decrease) and for *D. plantarius* (66% decrease; figure 3). Both species should be able to fill the suitable range towards 2070, but both should have a limited spread on the range of suitable habitat under Disp (figure 3 and 4; 14% increase under BLU and 4% under Disp for *D. fimbriatus*; 161% and 16%, respectively, for *D. plantarius*). The range of both species should shrink under DispCS (81% in 2050 and 76% in 2070, compared to current suitable habitat for *D. fimbriatus*; 88% and 53%, respectively, for *D. plantarius*).

**Figure 4:**
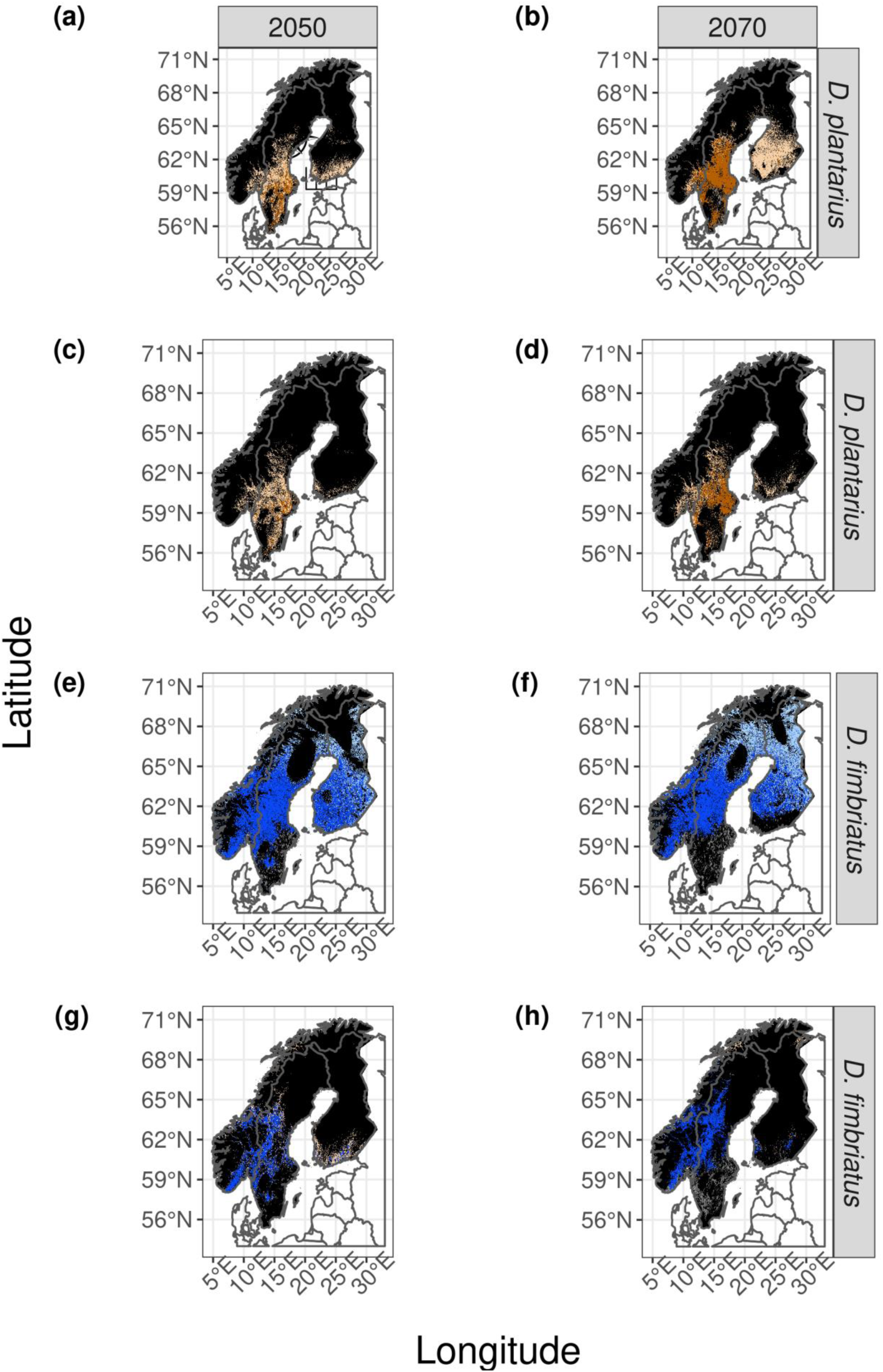
Map of the forecasted suitable habitat with an estimation of the reachable range predicted by the dispersion model (Disp) and reachable area from the connectivity model (DispCS) under the RCP4.5 scenario (RCP: Representative Concentration Pathway; in dark brown the reachable habitat for *D. plantarius* under Disp (a and b) and DispCS (c and d); in dark blue the reachable for *D. fimbriatus* under Disp (e and f) and DispCS (g and h); in black: unsuitable habitat; in grey: previously occupied habitat lost; in light brown and light blue: suitable but non reachable habitat).

The southern part of the suitable range should shrink, especially in Sweden and, to a lesser extent, in Finland. This range should expand in northern Fennoscandia (figure 4). According to model dispCS, tis shift should occur towards the North-Est, with a limited spread in southern Finland (figure 3). Similarly, the range of suitable habitat for *D. plantarius* should also increase towards the North-East under model Disp (figure 5). The shift of the centre of gravity is at a higher distance for the models which exclude Dispersal (Bioc and BLU) than model including dispersal (Disp and DispCS). The centre of gravity shifts farther without dispersal (models Bioc and BLU) than with dispersal (models Disp and DispCS).

**Figure 5:**
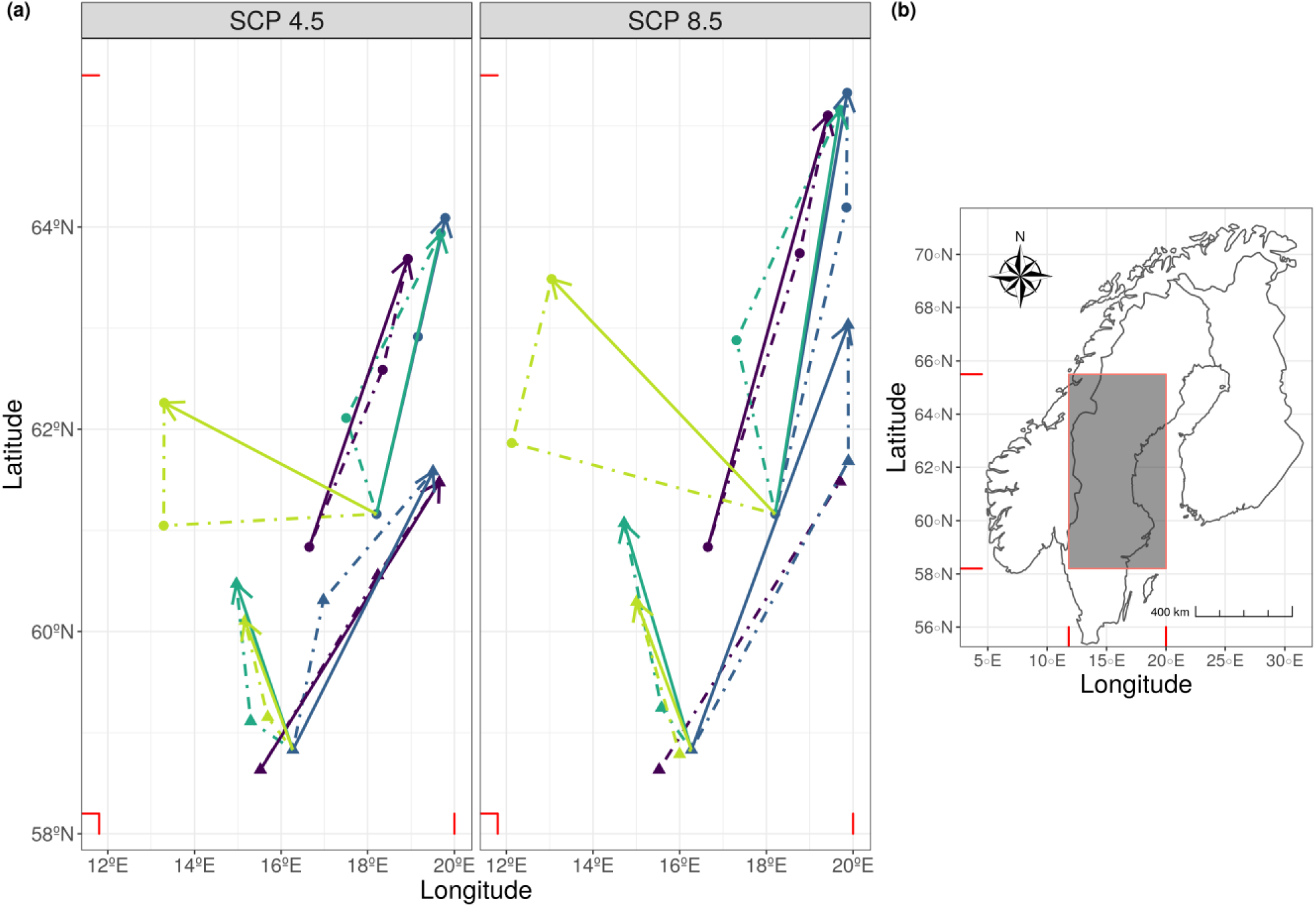
Shift in the centre of gravity of the two species distributions predicted by the four SDMs; solid lines: shift from current to 2070; dashed lines: shift from current time to 2050 and from 2050 to 2070. Dark purple: Bioc model; dark blue: BLU model; turquoise: Disp model; green: DispCS model.

The predicted distribution overlap between species was higher when considering only climatic variables than when accounting for land use at current time (Bioc model). Under the BLU model, the overlap should increase through time and is more important for the scenario SRCRCP8.5 than the 4.5 one (Schoener’s D values ranging from 0.55 at current time to 0.62 in 2070 for RCP4.5, it reached 0.68 under 8.5). The overlap should mainly occur at the Southern range of *Dolomedes fimbriatus* distribution (figure 6; Supplementary material Appendix 4, Tab. A4).

**Figure 6:**
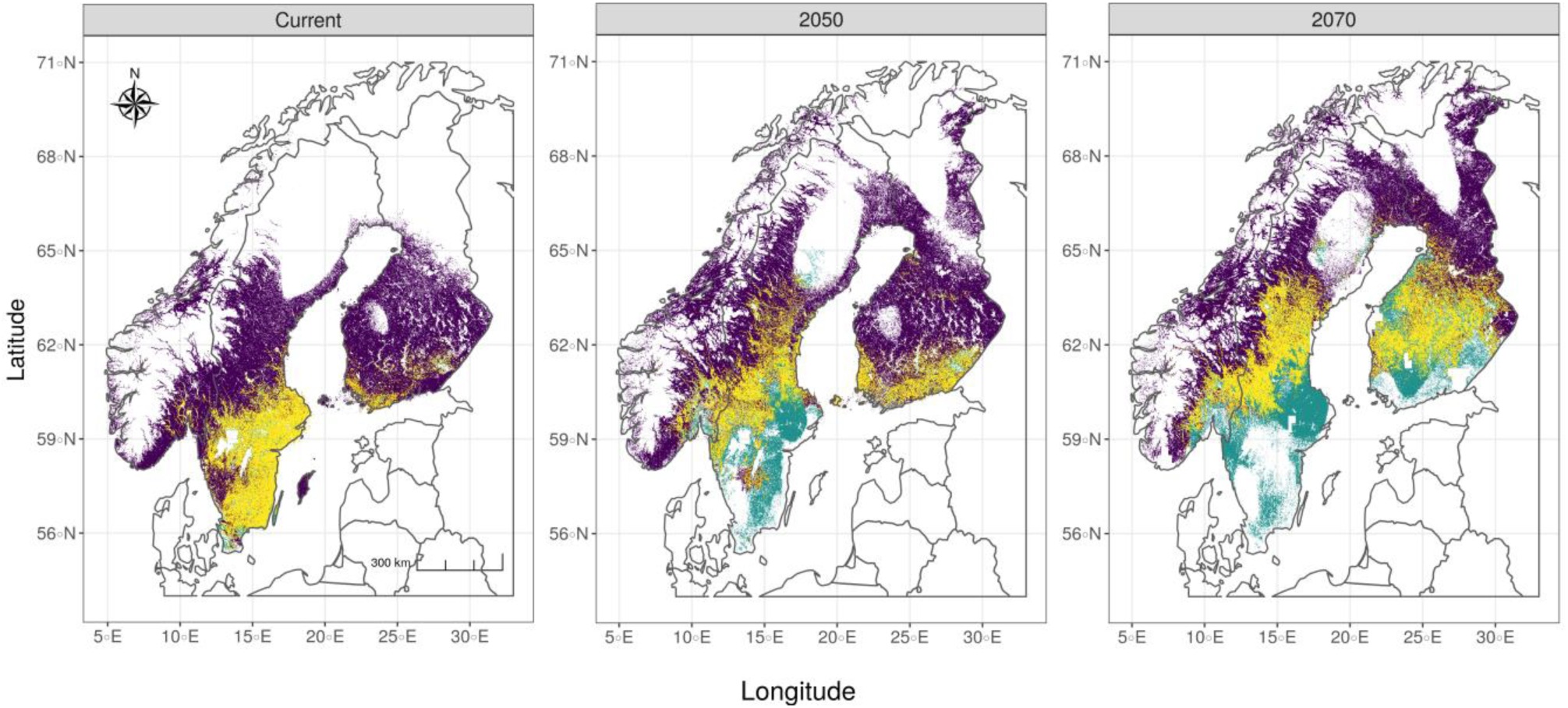
Range overlap predicted by model BLU from current time to 2070 under scenario RCP4.5. In addition to overlap of suitable range, suitable habitat for each species is represented.dark purple: suitable habitat for *D. fimbriatus*; green: suitable habitat for *D. plantarius*; yellow: suitable habitat overlapping between the 2 species.

## Discussion

Using species distribution models (SDM), we highlighted different range expansions and shifts of two closely related fishing spiders species in Fennoscandia. According to our predictions, the range of suitable habitat should spread for both D. fimbriatus and D. plantarius. Our climatic and habitat models (Bioc and BLU) confirmed the expansion of *D. plantarius* in Fennoscandia predicted by Leroy et al. (2013, 2014). In contrast, our hybrid models including dispersal and landscape connectivity (Disp and DispCS) predicted a more limited expansion.

### Northward range expansion of both *Dolomedes* species

A northward expansion in Fennoscandia is expected for the two species under both Bioc and BLU models. The range of suitable habitat should increase with the intensity of the climate change for *D. plantarius* and for *D. fimbriatus* in 2050. This northward expansion s also predicted in other taxa, as climate change promote an expansion of the range at the colder margin (Parmesan and Yohe 2003, Parmesan 2006). An increase in annual mean temperature and in temperature of the warmest month, which are the most important variables for both models, could impact the lifespan of the two spider species, and affect their distribution. Higher temperatures could increase the suitable period to produce juveniles, which could in turn increase the number of juveniles dispersing. The temperature encountered by juveniles also influences the dispersal ability and mode (ie, long vs short distance dispersal; Bonte et al. 2008b). Moreover, latitude and climate affect the time at which the *Dolomedes* reach maturity (Duffey 2012). This could increase the frequency of a second brood, which we already observed in September (unpublished data). Such an increase in temperature could, in turn, influence the speed of colonization of new habitats. The inclusion of land use in BLU models shrinks the range of suitable habitat, which confirms results from other, similar studies (Eskildsen et al. 2013)

Under the Disp model, suitable habitat should be less reachable for *D. plantarius* than for *D. fimbriatus*. The size of the area reached under the Disp model should be smaller than the current area for both species. In 2070, *D. fimbriatus* should have a range slightly equivalent to the suitable habitat estimated under BLU, whereas it should be smaller for *D. plantarius.* The limited expansion o*f D. plantarius* is explained mainly by dispersal ability. Indeed, we observe fewer spiderlings of *D. plantarius* showing dispersal behaviours, including long-distance dispersal through ballooning (unpublished data). Non-filling the suitable range is usually explained by either or both past and current limited dispersal, as exemplified by trees species (Svenning and Skov 2004). Some species are trapped in their geographical range, species which distribution has not changed since the last glaciation. Under a changing climate, species answer whether through microevolution or adaptive phenotypic plasticity (Radchuk et al. 2019). Some species are not yet able to adjust their phenology and physiology to changes induced by climate change. The importance of short-distance dispersal in fishing spiders should nonetheless maintain genetic exchange, or avoid genetic drift, at a smaller scale (Bell et al. 2005). A possible prevalence of this behaviour might also reinforce the importance of shorter dispersal as climate change and other factors like the increase of habitat fragmentation decrease long-distance dispersal of spiders (Bonte et al. 2006).

### Geographic range overlap and coexistence

The geographic and climatic niche of *D. plantarius* are included in the realised niche of *D. fimbriatus*. The first is a habitat specialist, the last is a generalist species living in a wider variety of environmental conditions over its range. Climate change increases the chance of overlap between these two sister species. However, we did not make predictions at a meso- or microhabitat scale, which would be too fine for SDMs. Yet, field observations suggest that both Dolomedes species also co-occur at finer spatial scales (Duffey 2012). The discrete nature and propensity to hide and dive of *D. plantarius* (van Helsdingen 1993), together with possible misidentification (Bellvert et al. 2013, Ivanov et al. 2017) might explain the small number of records and of co-occurrences. In North America, closely related species of *Dolomedes* like *D. trition* and *D. vittatus* were reported to co-occur at small spatial scales (Carico 1973).

Usually, closely related species co-occur less often than moderately related species (Weinstein et al. 2017). One one hand, an increase in co-occurrence might limit the distribution by segregation at the landscape scale. Indeed, the number of interactions between species in the ecosystem increases with climate change (Montoya and Raffaelli 2010), which generates a spatial separation between generalist and specialist species. Sympatric sister species usually diverge ecologically (Losos 2008), *Dolomedes* species differ in terms of habitat use (Duffey 2012). *D. plantarius* needs open habitat with slow-flowing water and water all year, while these factors do not seem to restrict *D. fimbriatus* (unpublished data). On the other hand, spatial segregation might occur at the micro-habitat scale. For instance, a study on *Tetragnatha* spiders showed that one of two co-existing spider species builds nursery webs higher in the vegetation when they co-occur (Williams et al. 1995). Finally, an increase in co-occurrence might lead to phenological shift in co-existence sites. Our observation in two Swedish locations of *D. fimbriatus* females with juveniles in the nursery while *D. plantarius* still carried egg sacs could support this. Other closely related wolf spider species (Lycosidae) also show differences in the timing of their breeding season to avoid intraguild predation (Balfour et al. 2003).

### Intrinsic limits of hybrid SDMs

Ideally, a mechanistic model should account for all phases of dispersal, ie, emigration, transfer, settlement, (Travis et al. 2012, 2013). The SDM accounting for dispersal which we used here it not a mechanistic model but is rather based on assumptions concerning the three stages of passive dispersal. Further studies should consider factors which influence individuals’ dispersal such as food availability (Bonte et al. 2008a), presence of endosymbionts (Goodacre et al. 2009), presence of conspecific in the short-long distance dispersal allocation (De Meester and Bonte 2010), or genetically inherited boldness (Bonte and Lens 2007). Since dispersal is not homogeneous within and among species (Clobert et al. 2009), a more realistic model should include information on dispersal and population size for each presence observation. The sampling of all sites is necessary to collect this information. There is a considerable gap between the theory and actual applications of data-demanding mechanistic SDMs (Briscoe et al. 2019). Knowing that the most used habitat is not necessarily the most suitable for the fitness of the species (Titeux et al. 2019), we used a hybrid model based on the lack of sufficient data for a full mechanistic model.

Less snow cover leads to less insulation, and thus, to colder subnivean habitat, where fishing spiders overwinter (Slatyer et al. 2017). Accounting for thermal niche information is possible with mechanistic models (Ceia-Hasse et al. 2014; Sinervo et al. 2010). However, the current knowledge of eco-physiological responses of fishing spiders to climate change is too scarce to allow fully mechanistic models.

### Conservation of fishing spiders

Fennoscandia may become a climatic refugium for *D. plantarius* as its range in continental Europe is expected to decrease (Leroy et al. 2013, 2014). The stronger the climate is, the more likely Fennoscandia will act as a refugium. The overlap between the two *Dolomedes* species should also increase with the climate change intensity. Arthropods conservation is challenging because of the fine-grain level needed as compared to vertebrates, the low empathy towards invertebrates, and the lowest number of conservation specialists available (Cardoso et al. 2020, Samways et al. 2020). Nonetheless, spiders have already been used as bio-indicators (Marc et al. 1999, Prieto-Benítez and Méndez 2011). Our models suggest that the conservation of both species is necessary as the reachable range size should drastically decrease in the future when accounting for dispersal and landscape connectivity. Conservation of preserved sites in a stepping-stones scheme is an alternative for species that are not able to use corridors (Noss and Daly 2006). Maintaining interconnected suitable sites in the first five kilometres around sites with known presence should help conserve current sites and promote expansion. With respect to fishing spiders, priority should be given to sites in southern Finland and central Sweden, where there is limited connectivity, and the spread of *Dolomedes* species is limited. Since *D. fimbriatus* has higher dispersal abilities, improving the connectivity in the North of the suitable range to make it reachable should improve the future range.

This work, together with other studies on *Dolomedes*, could be used to update the now outdated range assessment of *D. plantarius* (World Conservation Monitoring Centre 1996). The species’ conservation would benefit from such an update.

## Acknowledgements

We thank Stefano Mammola for useful comments and discussions on an early version of the manuscript. We also thank all the landowners who gave access to their properties.

## Author contributions

All authors contributed to the design and implementation of the research. JM analysed the data and drafted the manuscript. All authors contributed to writing of the manuscript and approved of the final version.

## References

Allouche, O. et al. 2006. Assessing the accuracy of species distribution models: prevalence, kappa and the true skill statistic (TSS). - J. Appl. Ecol. 43: 1223–1232.

Araújo, M. B. and New, M. 2007. Ensemble forecasting of species distributions. - Trends Ecol. Evol. 22: 42–47.

Balfour, R. A. et al. 2003. Ontogenetic shifts in competitive interactions and intra-guild predation between two wolf spider species. - Ecol. Entomol. 28: 25–30.

Barbet-Massin, M. et al. 2012. Selecting pseudo-absences for species distribution models: how, where and how many? - Methods Ecol. Evol. 3: 327–338.

Bell, J. R. et al. 2005. Ballooning dispersal using silk: world fauna, phylogenies, genetics and models. - Bull. Entomol. Res. 95: 69–114.

Bellard, C. et al. 2012. Impacts of climate change on the future of biodiversity: Biodiversity and climate change. - Ecol. Lett. 15: 365–377.

Bellard, C. et al. 2013. Will climate change promote future invasions? - Glob. Change Biol. 19: 3740–3748.

Bellvert, A. et al. 2013. First record of Dolomedes plantarius (Clerck, 1758)(Araneae: Pisauridae) from the Iberian Peninsula. - Rev. Ibérica Aracnol. 23: 109–111.

Bocedi, G. et al. 2017. RangeShifter: a platform for modelling spatial eco-evolutionary dynamics and species’ responses to environmental changes. - Methods Ecol. Evol.: 388–396.

Bonte, D. and Lens, L. 2007. Heritability of spider ballooning motivation under different wind velocities. - Evol. Ecol. Res. 9: 817–827.

Bonte, D. et al. 2006. Geographical variation in wolf spider dispersal behaviour is related to landscape structure. - Anim. Behav. 72: 655–662.

Bonte, D. et al. 2008a. Starvation affects pre-dispersal behaviour of Erigone spiders. - Basic Appl. Ecol. 9: 308–315.

Bonte, D. et al. 2008b. Thermal conditions during juvenile development affect adult dispersal in a spider. - Proc. Natl. Acad. Sci. 105: 17000–17005.

Bonte, D. et al. 2009. Repeatability of dispersal behaviour in a common dwarf spider: evidence for different mechanisms behind short- and long-distance dispersal. - Ecol. Entomol. 34: 271–276.

Braunisch, V. et al. 2013. Selecting from correlated climate variables: a major source of uncertainty for predicting species distributions under climate change. - Ecography 36: 971–983.

Briscoe, N. J. et al. 2019. Forecasting species range dynamics with process-explicit models: matching methods to applications. - Ecol. Lett. 22: 1940–1956.

Buisson, L. et al. 2010. Uncertainty in ensemble forecasting of species distribution. - Glob. Change Biol. 16: 1145–1157.

Cardoso, P. et al. 2020. Scientists’ warning to humanity on insect extinctions. - Biol. Conserv. 242: 108426.

Carico, J. E. 1973. The Nearctic species of the genus *Dolomedes* (Araneae: Pisauridae). - Bull. Mus. Comp. Zool. Harv. Coll. 144: 435–488.

Ceia-Hasse, A. et al. 2014. Integrating ecophysiological models into species distribution projections of European reptile range shifts in response to climate change. - Ecography 37: 679–688.

Clobert, J. et al. 2009. Informed dispersal, heterogeneity in animal dispersal syndromes and the dynamics of spatially structured populations. - Ecol. Lett. 12: 197–209.

De Meester, N. and Bonte, D. 2010. Information use and density-dependent emigration in an agrobiont spider. - Behav. Ecol. 21: 992–998.

Dorman, C. et al. 2012. Collinearity: a review of methods to deal with it and a simulation study evaluating their performance. - Ecography 36: 27–46.

Duffey, E. 1995. The distribution, status and habitat of *Dolomedes fimbriatus* (Clerck) and *D. plantarius* (Clerck) in Europe. - Proc. 15th Eur. Colloq. Arachnol.: 54–65.

Duffey, E. 2012. *Dolomedes plantarius* (Clerck, 1757) (Araneae: Pisauridae): a reassessment of its ecology and distribution in Europe, with comments on its history at Redgrave and Lopham Fen, England. - Bull. Br. Arachnol. Soc. 15: 285–292.

EEA, 2018. European Union, Copernicus Land Monitoring Service 2018, European Environment Agency (EEA)

Engler, R. and Guisan, A. 2009. MigClim: Predicting plant distribution and dispersal in a changing climate. - Diversity and Distributions 15: 590–601.

Eskildsen, A. et al. 2013. Testing species distribution models across space and time: high latitude butterflies and recent warming. - Glob. Ecol. Biogeogr. 22: 1293–1303.

ESRI 2009. World Imagery.

Etherington, T. R. 2016. Least-Cost Modelling and Landscape Ecology: Concepts, Applications, and Opportunities. - Curr. Landsc. Ecol. Rep. 1: 40–53.

Fawcett, T. 2006. An introduction to ROC analysis. - Pattern Recognit. Lett. 27: 861–874.

Febbraro, M. D. et al. 2019. Integrating climate and land-use change scenarios in modelling the future spread of invasive squirrels in Italy. - Divers. Distrib. 25: 644–659.

Fick, S. E. and Hijmans, R. J. 2017. WorldClim 2: new 1-km spatial resolution climate surfaces for global land areas. - Int. J. Climatol. 37: 4302–4315.

Finlayson, C. M. et al. 2019. The Second Warning to Humanity – Providing a Context for Wetland Management and Policy. - Wetlands 39: 1–5.

Garcia, R. A. et al. 2014. Multiple Dimensions of Climate Change and Their Implications for Biodiversity. - Science 344: 1247579.

GBIF: The Global Biodiversity Information Facility 2019. What is GBIF? - https://www.gbif.org/what-is-gbif

Goodacre, S. L. et al. 2009. Microbial modification of host long-distance dispersal capacity. - BMC Biol. 7: 32.

Grenouillet, G. et al. 2011. Ensemble modelling of species distribution: the effects of geographical and environmental ranges. - Ecography 34: 9–17.

Guisan, A. and Thuiller, W. 2005. Predicting species distribution: offering more than simple habitat models. - Ecol. Lett. 8: 993–1009.

Guisan, A. et al. 2013. Predicting species distributions for conservation decisions. - Ecol. Lett. 16: 1424–1435.

Hao, T. et al. 2019. A review of evidence about use and performance of species distribution modelling ensembles like BIOMOD. - Divers. Distrib. 25: 839–852.

Hijmans, R. J. and Graham, C. H. 2006. The ability of climate envelope models to predict the effect of climate change on species distributions. - Glob. Change Biol. 12: 2272–2281.

Hijmans, R. J. et al. 2005. Very high resolution interpolated climated surfaces for global land areas. - International Journal of Climatology 25: 1965–1978.

Hill, J. K. et al. 1999. Evolution of flight morphology in a butterfly that has recently expanded its geographic range. - Oecologia 121: 165–170.

Holec, M. 2000. Spiders (Araneae) of the fishpond eulittoral zone. - Ekológica Bratisl. 19: 51–54.

Hurtt, G. C. et al. 2011. Harmonization of land-use scenarios for the period 1500–2100: 600 years of global gridded annual land-use transitions, wood harvest, and resulting secondary lands. - Clim. Change 109: 117.

IPCC 2007. Climate Change 2007: The Physical Science Basis. Contribution of Working Group I to the Fourth Assessment Report of the Intergovernmental Panel on Climate Change.

Ivanov, V. et al. 2017. *Dolomedes Plantarius* (Araneae, Pisauridae) in Belarus: Records, Distribution and Implications for Conservation. - Arachnol. Mitteilungen 54: 33–37.

Kearney, M. 2006. Habitat, Environment and Niche: What Are We Modelling? - Oikos 115: 186–191.

Kearney, M. and Porter, W. 2009. Mechanistic Niche Modelling: Combining Physiological and Spatial Data to Predict Species’ Ranges. - Ecology Letters 12: 334–350.

Keeley, A. T. H. et al. 2017. Habitat suitability is a poor proxy for landscape connectivity during dispersal and mating movements. - Landsc. Urban Plan. 161: 90–102.

Keppel, G. and Wardell-Johnson, G. W. 2012. Refugia: keys to climate change management. - Glob. Change Biol. 18: 2389–2391.

Lafage, D. and Pétillon, J. 2016. Relative importance of management and natural flooding on spider, carabid and plant assemblages in extensively used grasslands along the Loire. - Basic Appl. Ecol. 17: 535–545.

Lafage, D. et al. 2015. Disentangling the influence of local and landscape factors on alpha and beta diversities: opposite response of plants and ground-dwelling arthropods in wet meadows. - Ecol. Res. 30: 1025–1035.

Lee, V. M. J. et al. 2015. Ballooning behavior in the golden orbweb spider *Nephila pilipes* (Araneae: Nephilidae). - Front. Ecol. Evol. in press.

Leroy, B. et al. 2013. First Assessment of Effects of Global Change on Threatened Spiders: Potential Impacts on *Dolomedes Plantarius* (Clerck) and Its Conservation Plans. - Biol. Conserv. 161: 155–163.

Leroy, B. et al. 2014. Forecasted climate and land use changes, and protected areas: the contrasting case of spiders. - Divers. Distrib. 20: 686–697.

Losos, J. B. 2008. Phylogenetic niche conservatism, phylogenetic signal and the relationship between phylogenetic relatedness and ecological similarity among species. - Ecol. Lett. 11: 995–1003.

Mammola, S. and Isaia, M. 2017. Rapid poleward distributional shifts in the European cavedwelling Meta spiders under the influence of competition dynamics. - J. Biogeogr. 44: 2789–2797.

Marc, P. et al. 1999. Spiders (Araneae) useful for pest limitation and bioindication. - Agric. Ecosyst. Environ. 74: 229–273.

McRae, B. H. et al. 2008. Using Circuit Theory to Model Connectivity in Ecology, Evolution, and Conservation. - Ecology 89: 2712–2724.

Melo-Merino, S. M. et al. 2020. Ecological niche models and species distribution models in marine environments: A literature review and spatial analysis of evidence. - Ecological Modelling 415: 108837.

Merow, C. et al. 2011. Developing Dynamic Mechanistic Species Distribution Models: Predicting Bird-Mediated Spread of Invasive Plants across Northeastern North America. - Am. Nat. 178: 30–43.

Miller, J. 2010. Species Distribution Modeling. - Geogr. Compass 4: 490–509.

Montoya, J. M. and Raffaelli, D. 2010. Climate change, biotic interactions and ecosystem services. - Philos. Trans. R. Soc. B Biol. Sci. 365: 2013–2018.

Noss, R. F. and Daly, K. M. 2006. Incorporating connectivity into broad-scale conservation planning. - In: Crooks, K. R. and Sanjayan, M. E. (eds), Connectivity conservation. Conservation biology. Cambridge University Press, pp. 587–619.

Parmesan, C. 2006. Ecological and Evolutionary Responses to Recent Climate Change. - Annu. Rev. Ecol. Evol. Syst. 37: 637–669.

Parmesan, C. and Yohe, G. 2003. A globally coherent fingerprint of climate change impacts across natural systems. - Nature in press.

Pelletier, D. et al. 2014. Applying Circuit Theory for Corridor Expansion and Management at Regional Scales: Tiling, Pinch Points, and Omnidirectional Connectivity. - PLoS One 9: e84135.

Pereira, H. M. et al. 2010. Scenarios for Global Biodiversity in the 21st Century. - Science 330: 1496–1501.

Prieto-Benítez, S. and Méndez, M. 2011. Effects of land management on the abundance and richness of spiders (Araneae): A meta-analysis. - Biol. Conserv. 144: 683–691.

Qiao, H. et al. 2015. No silver bullets in correlative ecological niche modelling: insights from testing among many potential algorithms for niche estimation. - Methods Ecol. Evol. 6: 1126–1136.

R Core Team 2019. R: A language and environment for statistical computing.

Radchuk, V. et al. 2019. Adaptive responses of animals to climate change are most likely insufficient. - Nat. Commun. 10: 1–14.

Reynolds, A. M. et al. 2007. Ballooning dispersal in arthropod taxa: conditions at take-off. - Biol. Lett. 3: 237–240.

Richmond, O. M. W. et al. 2010. Is the Climate Right for Pleistocene Rewilding? Using Species Distribution Models to Extrapolate Climatic Suitability for Mammals across Continents. - PLoS One in press.

Rissler, L. J. 2016. Union of Phylogeography and Landscape Genetics. - Proc. Natl. Acad. Sci. U. S. A. 113: 8079–8086.

Rödder, D. and Engler, J. O. 2011. Quantitative metrics of overlaps in Grinnellian niches: advances and possible drawbacks. - Glob. Ecol. Biogeogr. 20: 915–927.

Samways, M. J. et al. 2020. Solutions for humanity on how to conserve insects. - Biol. Conserv. 242: 108427.

Senay, S. D. et al. 2013. Novel three-step pseudo-absence selection technique for improved species distribution modelling. - PLoS One in press.

Sinervo, B. et al. 2010. Erosion of Lizard Diversity by Climate Change and Altered Thermal Niches. - Science 328: 894–899.

Shah, V. B. and McRae, B. 2008. Circuitscape: a tool for landscape ecology. - Proc. 7th Python Sci. Conf. 7: 62–66.

Slatyer, R. A. et al. 2017. Measuring the effects of reduced snow cover on Australia’s alpine arthropods. - Austral Ecology 42: 844–857.

Soberon, J. and Peterson, A. T. 2005. Interpretation of Models of Fundamental Ecological Niches and Species’ Distributional Areas. - Biodivers. Inform. in press.

Storfer, A. et al. 2007. Putting the ‘Landscape’ in Landscape Genetics. - Heredity 98: 128–142.

Svenning, J.-C. and Skov, F. 2004. Limited filling of the potential range in European tree species. - Ecol. Lett. 7: 565–573.

Thomas, C. F. G. et al. 2003. Aerial activity of linyphiid spiders: modelling dispersal distances from meteorology and behaviour. - J. Appl. Ecol. 40: 912–927.

Thuiller, W. 2004. Patterns and uncertainties of species’ range shifts under climate change. - Glob. Change Biol. 10: 2020–2027.

Thuiller, W. et al. 2009. BIOMOD – a Platform for Ensemble Forecasting of Species Distributions. - Ecography 32: 369–373.

Thuiller, W. et al. 2019. Uncertainty in ensembles of global biodiversity scenarios. - Nat. Commun. 10: 1–9.

Titeux, N. et al. 2016. Biodiversity scenarios neglect future land-use changes. - Glob. Change Biol. 22: 2505–2515.

Titeux, N. et al. 2019. Ecological traps and species distribution models: a challenge for prioritizing areas of conservation importance. - Ecography in press.

Travis, J. M. J. et al. 2012. Modelling dispersal: an eco-evolutionary framework incorporating emigration, movement, settlement behaviour and the multiple costs involved. - Methods Ecol. Evol. 3: 628–641.

Travis, J. M. J. et al. 2013. Dispersal and species’ responses to climate change. - Oikos 122: 1532–1540.

van Helsdingen, P. J. 1993. Ecology and Distribution of Dolomedes in Europe (Araneida: Dolomedidae). - Boll Acc Gioenia Sci Nat 26: 181–187.

van Vuuren, D. P. et al. 2011. The representative concentration pathways: an overview. - Clim. Change 109: 5.

Wagner, H. H. and Fortin, M.-J. 2013. A conceptual framework for the spatial analysis of landscape genetic data. - Conserv. Genet. 14: 253–261.

Warren, D. L. et al. 2008. Environmental Niche Equivalency versus Conservatism: Quantitative Approaches to Niche Evolution. - Evol. Int. J. Org. Evol. 62: 2868–2883.

Weinstein, B. G. et al. 2017. The role of environment, dispersal and competition in explaining reduced co-occurrence among related species. - PLoS One 12: e0185493.

Williams, D. D. et al. 1995. Trophic dynamics of two sympatric species of riparian spider (Araneae: Tetragnathidae). - Can. J. Zool. 73: 1545–1553.

World Conservation Monitoring Centre 1996. The IUCN Red List of Threatened Species 1996. in press.

