## Supplemental material for "Explicit integration of dispersal-related metrics improves predictions of SDM in predatory arthropods"

### Appendix 1. References of the R packages used.

Appelhans, T. et al. 2017. mapedit: Interactive Editing of Spatial Data in R.

Chamberlain, S. and Boettiger, C. 2017. R Python, and Ruby clients for GBIF species occurrence data. - PeerJ PrePrints in press.

Cheng, J. et al. 2018. leaflet: Create Interactive Web Maps with the JavaScript “Leaflet” Library.

Engler, R. et al. 2013. MigClim: Implementing Dispersal into Species Distribution Models.

Freeman, E. A. and Moisen, G. 2008. PresenceAbsence: An R Package for Presence Absence Analysis. - Journal of Statistical Software 23: 1–31.

Thuiller, W. et al. 2016. biomod2: Ensemble Platform for Species Distribution Modeling.

VanDerWal, J. et al. 2019. SDMTools: Species Distribution Modelling Tools: Tools for processing data associated with species distribution modelling exercises.

Warren, D. et al. 2017. ENMTools: Analysis of niche evolution using niche and distribution models.

Wickham, H. et al. 2019. Welcome to the tidyverse. - Journal of Open Source Software 4: 1686.

Zizka, A. et al. 2019. CoordinateCleaner: standardized cleaning of occurrence records from biological collection databases. - Methods in Ecology and Evolution 10: 744–751.

#### Appendix 2. References for the GBIF occurrence data.

Blindheim T (2020). BioFokus. Version 1.1286. BioFokus. Occurrence dataset https://doi.org/10.15468/jxbhqx accessed via GBIF.org on 2020-02-03.. Accessed from R via rgbif (https://github.com/ropensci/rgbif) on 2020-02-03

Creuwels J (2020). Naturalis Biodiversity Center (NL) - Chelicerata and Myriapoda. Naturalis Biodiversity Center. Occurrence dataset https://doi.org/10.15468/63lozv accessed via GBIF.org on 2020-02-03.. Accessed from R via rgbif (https://github.com/ropensci/rgbif) on 2020-02-03

GBIF.org on 2020-02-03.. Accessed from R via rgbif (https://github.com/ropensci/rgbif) on 2020-02-03

Hårsaker K, Finstad A G (2020). Terrestrial and liminic invertebrates systematic collection NTNU University Museum. Version 1.253. NTNU University Museum. Occurrence dataset https://doi.org/10.15468/fsreqb accessed via GBIF.org on 2020-02-03.. Accessed from R via rgbif (https://github.com/ropensci/rgbif) on 2020-02-03

iNaturalist.org (2020). iNaturalist Research-grade Observations. Occurrence dataset https://doi.org/10.15468/ab3s5x accessed via GBIF.org on 2020-02-03.. Accessed from R via rgbif (https://github.com/ropensci/rgbif) on 2020-02-03

Kaitila J (2019). Finnish Entomological Database. Version 1.3. Finnish Biodiversity Information Facility. Occurrence dataset https://doi.org/10.15468/jlud8r accessed via GBIF.org on 2020-02-03.. Accessed from R via rgbif (https://github.com/ropensci/rgbif) on 2020-02-03

Lahti T (2017). Hatikka Observation Database. Version 1.1. Finnish Biodiversity Information Facility. Occurrence dataset https://doi.org/10.15468/te1t6l accessed via naturgucker.de. naturgucker. Occurrence dataset https://doi.org/10.15468/uc1apo accessed via GBIF.org on 2020-02-03.. Accessed from R via rgbif (https://github.com/ropensci/rgbif) on 2020-02-03

Norwegian Biodiversity Information Centre ., Hoem S (2020). Norwegian Biodiversity Information Centre - Other datasets. Version 13.117. The Norwegian Biodiversity Information Centre (NBIC). Occurrence dataset https://doi.org/10.15468/tm56sc accessed via GBIF.org on 2020-02-03. Accessed from R via rgbif (https://github.com/ropensci/rgbif) on 2020-02-03

Shah M, Coulson S (2020). Artportalen (Swedish Species Observation System). Version 92.177. ArtDatabanken. Occurrence dataset https://doi.org/10.15468/kllkyl accessed via GBIF.org on 2020-02-03.. Accessed from R via rgbif (https://github.com/ropensci/rgbif) on 2020-02-03

Telenius A, Ekström J (2020). Lund Museum of Zoology (MZLU). GBIF-Sweden. Occurrence dataset https://doi.org/10.15468/mw39rb accessed via GBIF.org on 2020-02-03.. Accessed from R via rgbif (https://github.com/ropensci/rgbif) on 2020-02-03

The International Barcode of Life Consortium (2016). International Barcode of Life project (iBOL). Occurrence dataset https://doi.org/10.15468/inygc6 accessed via GBIF.org on 2020-02-03.. Accessed from R via rgbif (https://github.com/ropensci/rgbif) on 2020-02-03

The Norwegian Biodiversity Information Centre ., Hoem S (2020). Norwegian Species Observation Service. Version 1.77. The Norwegian Biodiversity Information Centre (NBIC). Occurrence dataset https://doi.org/10.15468/zjbzel accessed via GBIF.org on 2020-02-03.. Accessed from R via rgbif (https://github.com/ropensci/rgbif) on 2020-02-03

### Appendix 3. Additional methodological details for Disp and DispCS

#### Method used to create landscape connectivity maps.

The resistance map was retrieved from the suitability prediction map. The transformation method was a negative exponential (equation 1) function from Keeley (2007):

$$(1)100 - 99 \times \frac{(1 - e^{-c \times s})}{1 - e^{-c}}$$

In this function, *c* is a constant and *s* the habitat suitability value in a pixel. We used a value for *c* which is 0.25 for an almost linear relation between suitability and resistance. For this value, when the suitability is 0/1000 the resistance value is the opposite 1000/0.

As the distribution will supposedly change through time, we didn't use the node-to-node method. The nodes' method would be the occurrences at the present time and do not change with the model. The method developed by Pelletier (2014) and implemented in Febbraro (2019) counteracts this short come. Indeed, it uses a “wall-to-wall” method. The conductivity of the landscape is estimated from one side to the other side of the tile in North-South and East-West. The tiles are reassembled, and the two directional current mosaic are multiplied to obtain the conductivity map.


Febbraro, M. D. et al. 2019. Integrating climate and land-use change scenarios in modelling the future spread of invasive squirrels in Italy. - Divers. Distrib. 25: 644–659.

Keeley, A. T. H. et al. 2017. Habitat suitability is a poor proxy for landscape connectivity during dispersal and mating movements. - Landsc. Urban Plan. 161: 90–102.

Pelletier, D. et al. 2014. Applying Circuit Theory for Corridor Expansion and Management at Regional Scales: Tiling, Pinch Points, and Omnidirectional Connectivity. - PLoS One 9: e84135.

Table A1: MigClim parameters used to include dispersion in SDM predictions.

| Parameters | MigClim parameter | D. plantarius | D. fimbriatus | Explanation |
|---|---|---|---|---|
| Dispersal events per year | [DispSteps] | 1 per year | 1 per year | At least one nursery is produced per female |
| Short distance dispersal | [dispKernel] | 59% probability that spiders use short distance dispersions (first five km) | 76.6% * probability that spiders use short distance dispersion | The rappelling in the model is illustrated by the probability to disperse to the nearby cell (from 1 to 5 km here) |
| Dispersal barrier and filter | [barrier] and [barrierType] | Imperviousness | Imperviousness | Spiders can't establish population on impervious ground (threshold: 20%) |
| Long Distance dispersal (LDD) | [lddFreq] | 0.14% | 0.029% | Proportion of spiders using ballooning |
| Minimum LDD | [lddMinDist] | 1 km | 6 km | The minimum distance of long distance dispersion |
| Maximum LDD | [lddMaxDist] | 500 km | 500 km | The maximum distance of long distance event 1 |
| Probability to produce propagules | [propaguleProd] and [iniMatAge] | Half at 2 years | Half at 2 years | Corresponds to the time for a colonized cells to have mature dispersers, here to produce adults |

#### Appendix 4: Models evaluation and additional results.

Table A1: List of variables included in the different models (Bioc: SDM based on bioclimatic variables; BLU: SDM based on bioclimatic and land-use variables; Disp: SDM based on bioclimatic, land-use and dispersal abilities; DispCS: based on bioclimatic, land-use, dispersal abilities and landscape connectivity).

| **Layer** | **Unit** | **Used for:** | **Mean (SD) current** |
|---|---|---|---|
| Annual mean temperature | °C | Bioc, BLU, Disp, DiscpCS | 1.63 (±2.87) |
| Mean diurnal range | °C | Bioc, BLU, Disp, DiscpCS | 7.66 (±1.06) |
| Mean temperature of the warmest month | °C | Bioc, BLU, Disp, DiscpCS | 17.88 (±2.85) |
| Mean temperature of the wettest quarter | °C | Bioc, BLU, Disp, DiscpCS | 9.97 (±4.24) |
| Annual precipitation | mm | Bioc, BLU, Disp, DiscpCS | 728.58 (±359.32) |
| Grassland | % | BLU, Disp, DiscpCS | 5.89 (±13.58) |
| Wetness probability index | % | BLU, Disp, DiscpCS | 10.52 (±20.41) |
| Forest density | % | BLU, Disp, DiscpCS | 37.42 (±25.27) |
| Imperviousness | % | Disp, DiscpCS | 1.30±2.32) |

Table A2: Evaluation of the ensemble SDMs with ROC and TSS and variable importance (Bio01: annual mean temperature; Bio02: mean diurnal temperature range; Bio05: temperature maximum of the warmest month; Bio08: minimum temperature of the wettest quarter; Bio12: annual precipitation; Forest: tree cover density; Grass: grassland cover density; WAW: water and wetness).

| | | TSS | ROC | Bio01 | Bio02 | Bio05 | Bio08 | Bio12 | Forest | Grass | WAW |
|---|---|---|---|---|---|---|---|---|---|---|---|
| BIOC | *D. plantarius* | 0.872 | 0.976 | 0.654 | 0.042 | 0.112 | 0.081 | 0.025 | X | X | X |
| | *D. fimbriatus* | 0.812 | 0.958 | 0.786 | 0.072 | 0.331 | 0.038 | 0.059 | X | X | X |
| BLU | *D. plantarius* | 0.926 | 0.993 | 0.665 | 0.029 | 0.115 | 0.098 | 0.049 | 0.07 | 0.01 | 0.10 |
| | *D. fimbriatus* | 0.872 | 0.984 | 0.785 | 0.066 | 0.263 | 0.032 | 0.046 | 0.13 | 0.03 | 0.16 |

Table A3: Estimated range expansion/reduction from current time between species, models and scenarios.

| Species | Model | Scenario | Current – 2050 | Current – 2070 |
|---|---|---|---|---|
| *D. fimbriatus* | Bioc | RCP4.5 | 34 | 40 |
| | | RCP8.5 | 49 | 50 |
| | BLU | RCP4.5 | 10 | 14 |
| | | RCP8.5 | 20 | -5 |
| | Disp | RCP4.5 | -20 | 4 |
| | | RCP8.5 | -29 | -8 |
| | DispCS | RCP4.5 | -81 | -76 |
| | | RCP8.5 | -90 | -82 |
| *D. plantarius* | Bioc | RCP4.5 | 156 | 271 |
| | | RCP8.5 | 273 | 391 |
| | BLU | RCP4.5 | 50 | 161 |
| | | RCP8.5 | 141 | 173 |
| | Disp | RCP4.5 | -66 | 16 |
| | | RCP8.5 | -72 | 0 |
| | DispCS | RCP4.5 | -88 | -53 |
| | | RCP8.5 | -93 | -89 |

Table A4: Variation of Schoener's D overlap value through time between models and scenarios

| Model | Scenario | Current | 2050 | 2070 |
|---|---|---|---|---|
| Bioc | SRC 4.5 | 0.63 | 0.67 | 0.64 |
| | SRC 8.5 | 0.63 | 0.64 | 0.65 |
| BLU | SRC 4.5 | 0.55 | 0.65 | 0.62 |
| | SRC 8.5 | 0.55 | 0.64 | 0.68 |
